# The interplay between moult of flight feathers and fuelling conducted on the breeding grounds of the Great Snipe *Gallinago media* from the eastern European, lowland population

**DOI:** 10.1101/2024.01.23.576834

**Authors:** Marta Witkowska, Michał Korniluk, Pavel Pinchuk, Tomasz Tumiel, Natalia Karlionova, Włodzimierz Meissner

## Abstract

The Great Snipe as a long-distant migrant wintering in Africa, faces the challenge of accumulating sufficient energy reserves before the departure from European breeding grounds. Despite possible trade-offs in resource allocation, this species additionally initiates moult of flight feathers before southward migration. Here we discuss the strategy of flight feather moult and fuelling, exploring their scheduling, constrained by the timing of breeding and departure for female and male Great Snipes from the European lowland population. We found significant intrasexual differences in both moult initiation date and moult duration. Males start flight feather replacement more than two weeks earlier and moult faster compared to females. However, neither sex completed this process on breeding grounds before the migration, as late in the season all males and half of the females had suspended their primary moult, with the remaining females not moulting at all. Moult of secondaries occurred exceptionally in the studied population. We observed a non-linear energetic stores gain in the studied period, where both sexes maintained a stable and low body condition until the end of July, coinciding with the primary moulting period. Subsequently, there was an increase in body condition, of approximately 1% of the lean body mass per day, indicating a shift towards fuelling for migratory flight. The overlap between stages of maintaining a stable and low body condition and moulting suggests a resource allocation towards feather growth before initiating fuelling. Our study describes moult strategy in Great Snipe conducted on their breeding grounds, highlighting the intrasexual differences, likely resulting from different parental duties of males and females of this lekking species.

## Introduction

In the annual cycle of most bird species of the temperate zone, there are crucial life-history events distinguished, such as breeding, migration, and moulting. As all of those are considered energetically demanding processes, which require a large amount of available resources to be fulfilled, birds tend to avoid overlapping them, and by that evade possible trade-offs in energy allocation (Ricklefs 1996, Hemborg & Lundberg 1998, Stutchbury *et al*. 2011). The timing of breeding and migration is for many species environmentally constrained (Wingfield 2008), but there is usually some more flexibility in fitting moult into the annual cycle (McNamara *et al*. 1998, Helm & Gwinner 2006, Conklin *et al*. 2013), which is reflected in a large variety of moulting strategies in birds (Kjellén 1994). For small and medium-sized bird species annual replacement of worn feathers is crucial for their survival and performance in life-history events, due to their functions in a wide range of processes in birds, including but not limited to locomotion, thermoregulation and communication (Terrill & Shultz 2022). Migratory species, and especially those that conduct long-distance flights, should benefit from extending the time of moult, as the structural quality of feather increases with the time it takes for it to grow, which makes them more resistant to wear and could improve the migratory performance (Dawson *et al*. 2000a, Serra 2001a, Vágási *et al*. 2012). On the other hand, most bird species separate active moult and migration, as it compromises fuelling for flight (Swaddle & Witter 1997, Stutchbury *et al*. 2011), and the occurrence of the wing-gap impairs flight, additionally increasing energetical demands (Swaddle & Witter 1997).

Reviewing moult strategies in relation to migration reveale-d that most wader species that are long-distance migrants do not complete their primary moult on their breeding grounds before the departure (Remisiewicz 2011). Their most common strategies are to conduct moult on wintering grounds (Kjellén 1994). It is thought to be caused by a longer time available for moulting on wintering grounds, where migrants usually spend a larger portion of the year, with migratory fuelling being the main temporal limitation for the timing of moulting. Moreover, availability of sufficient food resources further constrains scheduling of moult, and for long-distance migratory waders such predicable and rich resources are usually present on their southern, non-breeding grounds (Remisiewicz 2011). In comparison, reproduction carried out on breeding grounds together with the preparation for the southward migratory flight constrains time and resources available for moulting to a greater extent. Therefore, in those few wader species whose strategy is to begin moult on breeding grounds, prior to southward migration, both processes of feather growth and fuelling for a flight might overlap, and since both are energetically costly, birds should compromise resource allocation between them (Lindström *et al*. 1994, Rubolini *et al*. 2002, Bonier *et al*. 2007).

The deposition of fat and protein stores as fuel for flight is a key part of migration, that affects the speed and distance of the flight (Alerstam & Lindström 1990). It is also considered to be a determinant of the whole migration speed, as it takes longer than the migratory flight itself (Lindström *et al*. 2019). Since faster fuelling imposes increased migration speed, long-distance migrants covering a large range in non-stop flights, ought to maximize fuel deposition rates, because arriving sooner to the destination is considered advantageous (Faaborq *et al*. 2010, Morrison *et al*. 2019). However, this process can be limited by ecological factors such as insufficient food availability, or birds’ physiological constraints (Lindström 2003). Moreover, other life events may project on fuel deposition, for example in species with females and males having different parental duties, strategies for fuelling may differ between sexes (Mazur *et al*. 2021).

The Great Snipe *Gallinago media* is a wader species known for its lekking behaviour and long-distance migration (Cramp & Simmons 1983). Unlike the majority of waders, the primary moult of adult females and males in this species is initiated on breeding grounds, followed by birds suspending it before the departure, and later completing the process in sub-Saharan Africa (Debayle *et al*. 2017). Studies on migration revealed that males of this species are able to cover a large portion of the migratory distance with one, nonstop flight, with a high speed achieved. After trans-Saharan flight birds stay in the Sahel zone for approximately three weeks before making a second flight further sought to their final wintering grounds (Lindström *et al*. 2016). Moreover, their migration follows a diel cycle, with birds considerably changing their flight altitude between day and night (Lindström *et al*. 2021). Unfortunately, we lack knowledge about similar details of migration strategy and other biological aspects in females, as there is an existing disproportion of research, with most studies focused on males’ biology, especially in case of the lowland population (Korniluk *et al*. 2020, Witkowska *et al*. 2022, 2023, Korniluk & Chylarecki 2023). The pre-migratory processes conducted before departure to wintering grounds, such as initiation of moult of flight feathers and fuelling for long-distance, southward migratory flight were not yet investigated both in females and males of this species. Studying those processes would improve our understanding of migration strategy and the completion of moult achieved in Africa. In this work, we aimed to describe the strategy of moulting flight feathers on breeding grounds and fuelling for southward migratory flight for adult Great Snipe of the Eastern European, lowland population. We investigated intrasexual differences in both processes and how they fit together on a temporal scale limited by breeding and departure for wintering grounds.

## Materials & Methods

### Fieldwork

Data were collected between the year 2000 and 2023 in two locations occupied by the lowland population of Great Snipe (Cramp & Simmons 1983): 1) in the floodplain meadow in the valley of Pripyat River near Turov, Gomel Region, Belarus (52° 05′ N, 27° 46′ E), which is an important breeding and stopover site for waders during both spring and autumn migrations (Meissner *et al*. 2011, Pinchuk *et al*. 2016) and 2) in NE Poland within a floodplains and post-peatland meadows. In both sites adult Great Snipes, older than their second calendar year, were trapped from June till the early September, corresponding to the late part of the breeding season and the period of preparation for migratory flight for the studied species (Cramp & Simmons 1983). In Belarus Snipes were trapped on the leks using mist nets. To reduce the disturbance of lekking birds, capture events on leks were limited to 4 hours and were separated by at least 5-day break. In Poland Great Snipes were searched at night with a thermal cameras in potential foraging sites (flooded and mown meadows) within a few kilometres of known leks. Individuals were then spotted with headlights and captured with a landing net attached to a 5 m carbon fibre pole. In all captured individuals we measured the total head length, bill length and tarsus length (measured with callipers to the nearest 0.1 mm), wing length (all measured with a ruler to the nearest 1 mm), and weight (measured with an electronic balance to the nearest 0.1 gram), according to standard procedure (Busse & Meissner 2015). For each individual we recorded moult formula of primaries and secondaries, representing the stage of moulting of each feather as proposed by (Ginn & Melville 1983). Following this method, each bird’s moult state was described as ten digits representing ten primaries and ten digits representing ten secondaries respectively, where old feathers were coded as 0; new, fully grown feathers were coded as 5, and scores 1 to 4 are intermediate stages of feather development. Later, birds were sexed based on body measurements according to Höglund et al., (1990). We did not assign sex to individuals with measurements lying in the intermediate range between females and males. Overall, our sample contained 79 females and 223 males.

### Statistical analysis

#### Body condition

Body condition can be interpreted in various ways, however, in this work, we approached it as an amount of energetic resources gathered by the individual. To quantify it we used two morphometric variables: 1) scaled mass index as proposed in (Peig & Green 2009), where the body mass of an individual is corrected for its structural size and 2) body mass of individuals. The body mass of an individual is commonly interpreted by researchers as a measure of energetic resources gathered by the individual (Labocha & Hayes 2012). Although it may be biased in species with variety in size between individuals, e.g. in species with strong body size sexual dimorphism, using body mass as a predictor of condition may be advantageous, due to its easy interpretation and comparison between studies. Great Snipes exhibit reversed sexual size dimorphism, with females being larger than males (Hoglund *et al*. 1990). To account for differences in body size between sexes, in this study, we used scaled mass index as our primary estimate for body condition in our analysis of factors affecting moult of a primary feather, as well as the process of fuelling before departure. Additionally, we provided descriptive results of the analysis of the fuelling process using body mass, with the full results presented in the Supplementary Online Materials. We also established lean body mass for each of the sexes separately, as the mean body mass of 10% of the lightest individuals.

#### Moult

In Great Snipe, moult of primaries progresses from the innermost primary towards the outermost primary and generally two or three primaries are being moulted at the same time. In this species moult can be suspended on breeding grounds and later resumed on wintering grounds (Debayle *et al*. 2017). Based on that knowledge, all birds were grouped into one of the three moult stage categories: 1) moult not started, where we assigned birds with all 10 primaries scored as 0; 2) active moult, where we assigned birds with a number of adjacent primaries actively growing (scored from 1 to 4) in characteristic gradual pattern; and 3) suspended moult, where we assigned birds with a number of fully grown primaries (scored as 5) with the rest of the adjacent primary feathers next in moulting sequence being old (scored as 0). We did not report any individuals with a completed moult, therefore we did not account for such moult stage category. We used Pearson’s χ^2^ test with p-values computed by Monte Carlo simulation based on the 2000 replicates test to analyse differences in the proportion of birds in three established moult stage categories in a given half-month period, from the first half of June up until the second half of August. Similarly, we used the same statistical test to investigate variation between both sexes among individuals with suspended moult regarding differences in the range of moult of primaries conducted before suspending this process. In that case the range of moult of primaries was described as number of the last, most outermost renewed primary. For each individual, we calculated the Percentage of Feather Mass Grown (later referred to as PFMG) following the method described by Underhill & Zucchini, (1988), based on moult score of each primary and its mass reported for this species (Meissner et al., 2018). We used the Underhill-Zucchini moult model (later referred to as UZ moult model), computed with the *moult* package (Erni *et al*. 2013) in R version 4.2.2 (R Core Team 2022), to estimate moult parameters, such as mean start date and mean end date of moult and their standard deviation, as well as the duration of moult. Due to the lack of birds with primary moult finished, we decided to fit the type 5 model (Underhill *et al*. 1990) to our data, using data on birds with not started moult and birds actively moulting, excluding birds with suspended moult. The date of capture of a particular individual was stated as a day of the season, with the 1^st^ of June established as the first day of the season. To determine differences in moult strategy between both sexes and to check for the effect of body condition on this process, we also modelled mean start date, mean end date, and the duration of moult as a function of the sex of an individual and scaled mass index, separately in single-predictor models, as well as a global model including both predictors together. Later all three models with sex and/or body condition estimation as predictors and a starting model containing no predictors were ranked using Akaike Information Criterion corrected for small sample size (AIC_c_) and Akaike weights (w_i_) (Burnham & Anderson 2004).

As the moult of the secondaries was observed in very limited number of individuals, constituting less than 1% of the whole data set, we were unable to perform any statistical analysis on those. Therefore we chose a descriptive approach to present results concerning this process undertaken by adult Great Snipes.

#### Fuelling

We considered the same period for analysing fuelling before the southward migratory flight, as we did for moulting, with the 1^st^ of June established as the first day of the season. Preliminary analysis revealed a non-linear change in scaled mass index in time, therefore we decided to use the Generalized Additive Model (GAM) (Hastie & Tibsgirani 1986) to analyse the process of fuelling before departure, where scaled mass index was used as a dependent variable. In the set of proposed models, we included a null model, with intercept as the sole factor, the global model with the day of the season as a smooth term, sex, and its interaction with the day of the season as independent variables. We also established two reduced, nested models: a model with the day of the season as a single independent variable, and a model with the day of the season as a smooth term and the sex of an individual as an independent variable. Simillar to the UZ moult models used to analyse primary moult, in case of GAM model selection was carried out based on the Akaike Information Criterion and Akaike weights. Model fit was also described using R^2^ and the percentage of explained deviance. The models were fit using *gamm4* package (Wood *et al*. 2020), with model selection performed using *MuMIn* package (Bartoń 2023) in R version 4.2.2 (R Core Team 2022).

## Results

During the studied period we captured 12 females and 41 males actively moulting their primaries, as well as 11 females and 24 males showing suspended moult of primary feathers. Only in 3 males we detected signs of secondary mount, with all of them having their moult suspended, where only the first, innermost secondary was renewed, together with 6 (2 males) or 7 (1 male) primaries. The proportion of individuals in a given moult stage category differed between six established half month periods during the studied period both in females (χ^2^ test, *χ^2^*= 36.71, *P* < 0.001) and males (χ^2^ test, *χ^2^* = 186.5, *P* < 0.001). The general pattern showed the increase in proportion of birds with active moult and later with suspended moult in time, combined with the proportion of birds with moult not started decreasing over the season (Fig. 1). In both sexes first birds with active moult were caught in the second half of June and were present in the studied site up until the second half of July. There was no difference in the proportion of birds in different moult stage categories between sexes in the first two established periods (the first and second half of June; Table 1). However, later in the season males had a larger proportion of birds in active and/or suspended moult and a lower proportion of birds with moult not started compared to females. The last males with no signs of moult were captured in the first half of July, whereas females with all old primaries were present until the end of the field study in the second half of August (Fig. 1).

**Fig. 1.**
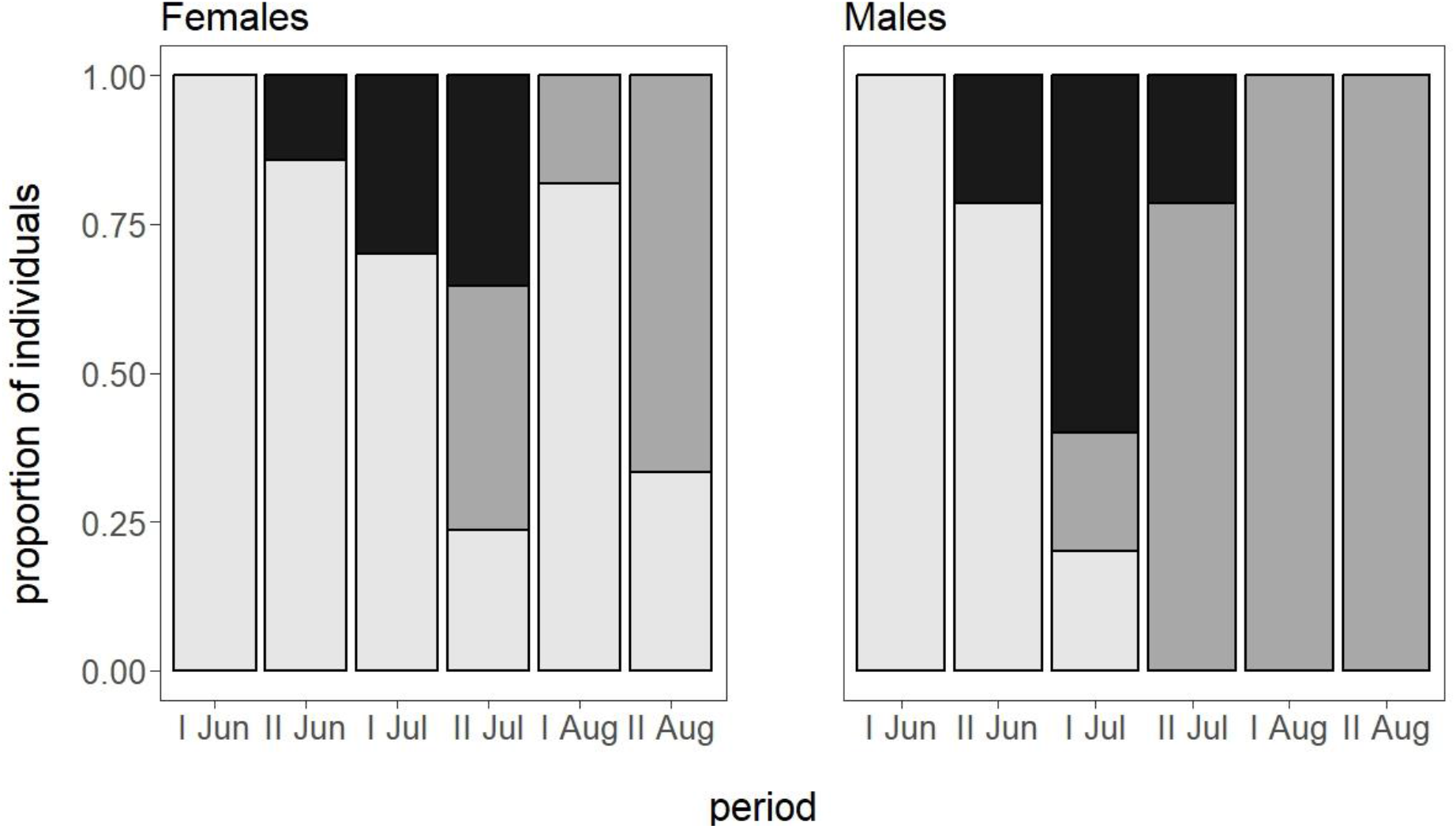
Proportion of individuals in adult females and males in three different moult stage categories (light grey – moult not started, dark grey – suspended moult, black – active moult) in six consecutive half-month periods.

**Fig. 2.**
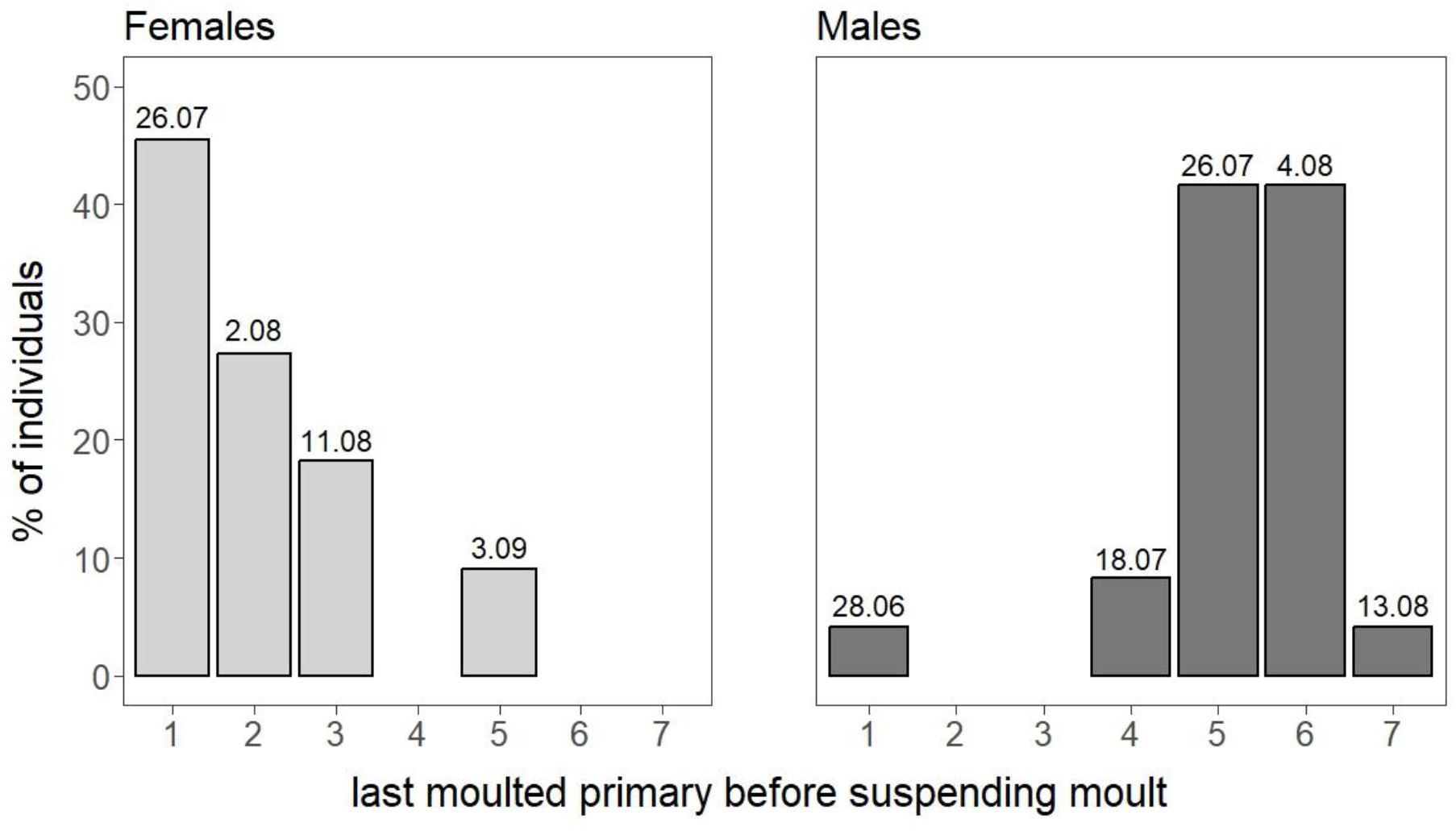
Percentage of individuals in adult females and males suspending their moult on a given primary. Date of completing moult of a given primary, estimated with UZ moult model type 5 is give above each bar.

**Table 1.** Comparison of frequencies of individuals in three moult categories between females and males in the half-month periods, based on the estimates of Pearson’s χ^2^ test with p-values computed by Monte Carlo simulation with the 2000 replicates. First half of June is excluded, due to only one level of variables (all birds were in the same moult stage category).

| Period | $\chi^2$ | *P* |
| --- | --- | --- |
| II June (16<sup>th</sup> June – 30<sup>th</sup> June) | 0.577 | 0.567 |
| I July (1<sup>st</sup> July – 15<sup>th</sup> July) | 6.875 | 0.024 |
| II July (16<sup>th</sup> July – 31<sup>st</sup> July) | 5.651 | 0.046 |
| I Aug (1<sup>st</sup> Aug – 15<sup>th</sup> Aug) | 11.455 | 0.001 |
| II Aug (16<sup>th</sup> Aug – 31<sup>st</sup> Aug) | 12.245 | <0.001 |

Females and males significantly differed in the acquired range of the suspended moult (χ^2^ test, *χ^2^* = 26.915, *P* < 0.001), where majority of females suspended their moult either on the 1^st^, innermost primary or on the 2^nd^ primary (respectably, 45% and 27% of all females with suspended moult). In males most individuals suspended their moult on the 5^th^ or 6^th^ primary (in both cases 48% of all males with suspended moult). No birds had moulted more than seven primaries before suspending their moult.

The UZ moult model with included sex of individuals as a factor influencing all three moult parameters had the best fit and was the most parsimonious among the set of proposed models, due to the lowest AICc values and the highest w_i_ values (model UZ1, Table 2). Therefore, presented moult parameters obtained with UZ moult models were estimated based on this model. Including scaled mass index in UZ moult models did not improve their fit (models UZ3 and UZ4, Table 2).

**Table 2.**
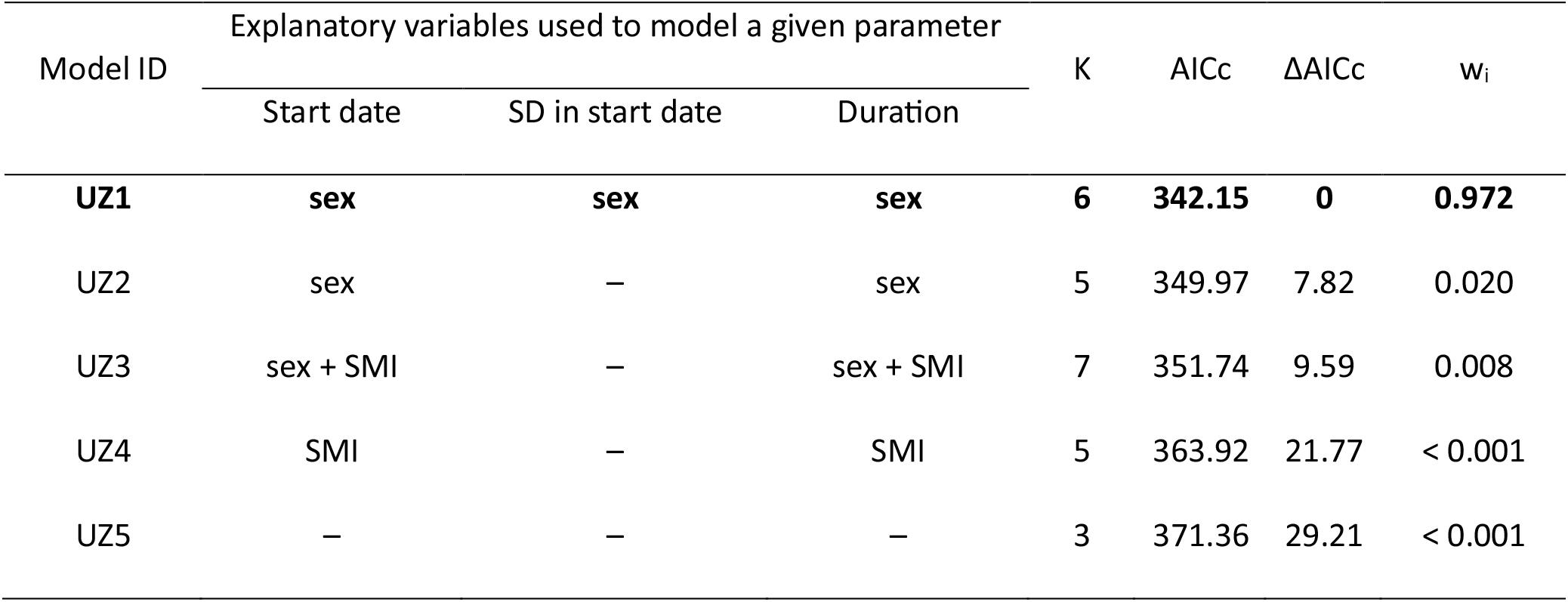
Ranking of UZ moult models used to estimate primary moult parameters for adult female and male Great Snipes. K – number of parameters, AICc – Akaike’s Information Criterion for small sample size, ΔAICc – difference in AICc between the given model and model with the lowest AICc value, w_i_ – Akaike weight. Top ranking model with the ΔAICc = 0 and the highest w_i_ is bolded.

According to the best fitted UZ moult model (model UZ1, Table 2) males started their primary moult in the end of June (mean start date 24^th^ June) and begin their primary moult on average 26 days earlier in comparison to females (Table 3). In females, the standard deviation of estimated start date was approximately 2.5 times larger than in males. Moreover, both sexes differed in primary moult duration, with males moulting their primaries approximately 1.4 times faster than females, resulting in their moult end date estimated to be on average 61 days earlier, assuming constant and uninterrupted progression of primary moult (Table 3, Fig. 3).

**Fig. 3.**
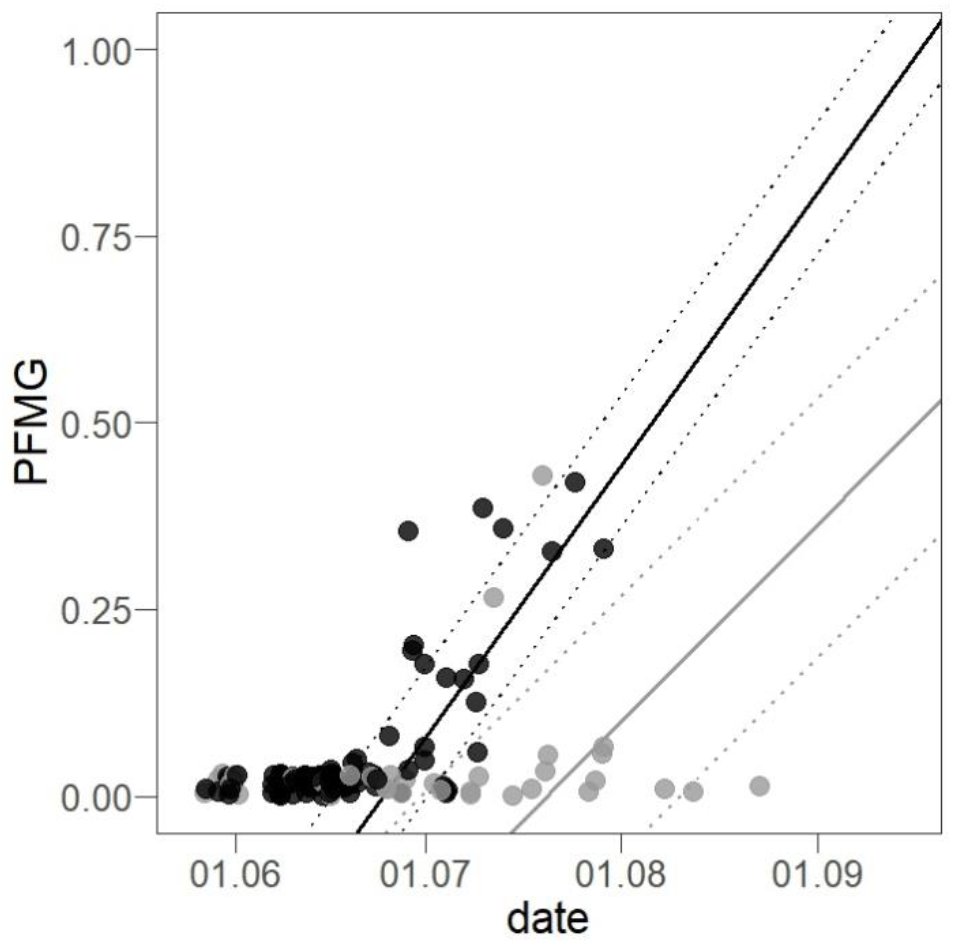
Changes of the percentage of feather mass grown (PFMG) in the season form females (grey) and males (black). Dots – data points representing individuals with a given PFMG in particular day of the season, line – progression of moult estimated with the type 5 UZ model, dotted line – standard deviation of the moult start date

Ranking of GAM models used to describe fuelling before departure revealed that the global model including day of the season as a smooth term, sex and its interaction with day of the season performed better than other proposed models (model SM1, Table 4). In this model we detected significant influence of the day of the season as a smooth term on scaled mass index and significant interaction between sex of an individual and the day of the season, where stable scaled mass index was detected until approximately 54^th^ day of the season (25^th^ July) with later increase. More rapid increase of this parameter was detected in males than in females (Fig. 4, Table 5). Both sexes however did not differ in overall scaled mass index (Table 5). Analysis including body mass instead of the scaled body mass index yielded similar results (Table 1S, Table 2S, Fig. 1S), however presenting it Supplementary Online Materials allows for the comparison between previous studies using this parameter.

**Fig. 4.**
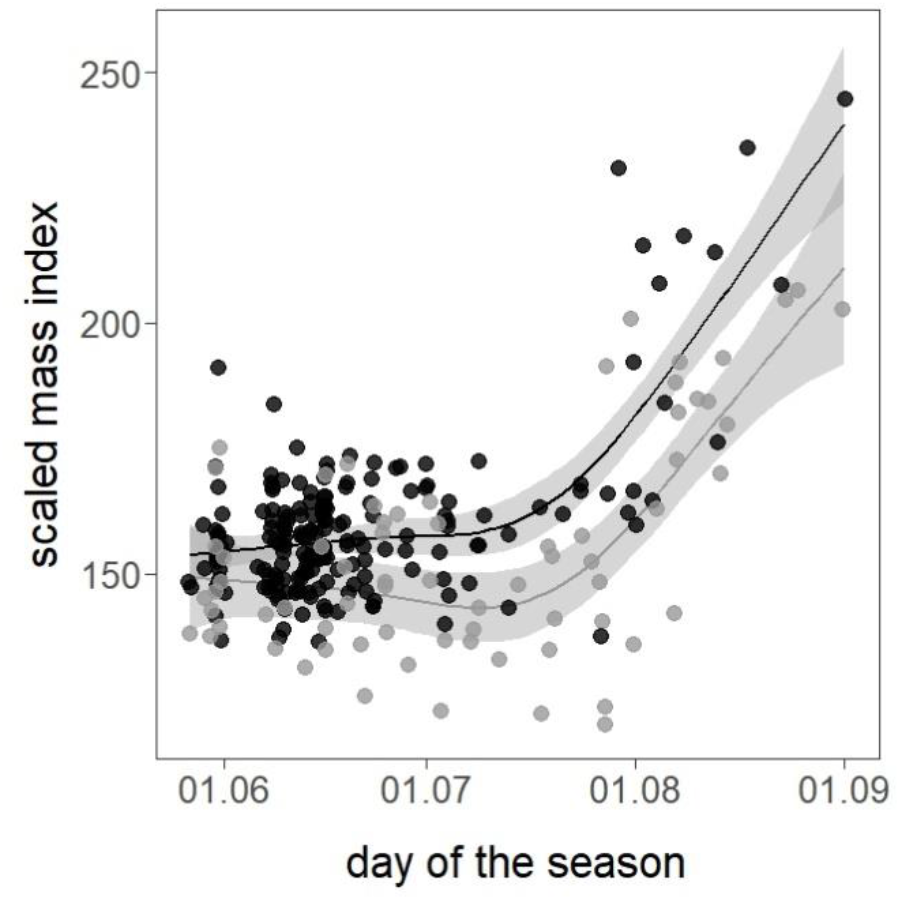
Changes of scaled mass index during the season in adult females (grey) and males (black). Line – significant relationship estimated with the Generalized Additive Model, grey area – 95% confidence interval.

**Table 3.** Primary moult parameters estimated with the top ranking type 5 UZ moult model (UZ1; Table 1) and sample size of birds in two moult stage categories included in the model.

| Sex | Moult Parameters |  |  |  |  |  |  |  | Sample size |  |
| --- | --- | --- | --- | --- | --- | --- | --- | --- | --- | --- |
|  | Start Date |  | SD in start date |  | Duration in days |  | End Date |  | Moult stage category |  |
|  | Estimate | SE | Estimate | SE | Estimate | SE | Estimate | SE | Moult not started | Active moult |
| Females | 20<sup>th</sup> Jul | 7.28 | 21.21 | 7.22 | 117.01 | 45.41 | 14<sup>th</sup> Nov | 7.28 | 50 | 12 |
| Males | 24<sup>th</sup> Jun | 7.38 | 8.45 | 1.02 | 85.04 | 46.41 | 17<sup>th</sup> Sep | 7.38 | 138 | 41 |

**Table 4.** Ranking of GAMs estimating the relationship of scaled mass index with sex and day of the season. 1 in model formula indicates null model with only intercept included as an independent variable. s() – independent variable used as a smooth factor, + - additive relationship, * - interaction between variables, edf – effective degrees of freedom, AICc – Akaike’s Information Criterion for small sample size, ΔAICc – difference in AICc between the given model and model with the lowest AICc value, w_i_ – Akaike weight. Adjusted R^2^ and percentage of explained deviance is given. Top ranking models with the ΔAICc < 2 are bolded.

| Model ID | Model formula | edf | AICc | ΔAICc | $w_i$ | adj. $R^2$ | % Dev. Exp. |
| --- | --- | --- | --- | --- | --- | --- | --- |
| **SMI1 (global)** | **SMI = s(day) + sex + sex\*day** | **9.07** | **2101.19** | **0** | **0.987** | **0.529** | **54.2%** |
| SMI2 | SMI = s(day) + sex | 8.23 | 2109.89 | 8.72 | 0.013 | 0.512 | 52.3% |
| SMI3 | SMI = s(day) | 6.22 | 2383.57 | 282.38 | < 0.001 | 0.434 | 44.2% |
| SMI4 (null) | SMI = 1 | 2 | 2546.73 | 445.54 | < 0.001 | <0.001 | 0% |

**Table 5.** Results of top ranking GAM (SMI1) explaining the relationship between scaled mass index and sex, interaction between sex and the day of the season and day of the season as a smooth factor.

| Fixed effect | Estimate | SE | z | P |
| --- | --- | --- | --- | --- |
| Intercept | 47.57 | 3.87 | 12.29 | <0.001 |
| sex (M) | 4.00 | 3.27 | 1.23 | 0.22 |
| sex (M) * day | 0.25 | 0.08 | 3.27 | 0.001 |
| Smoothed term |  |  | *F* | *P* |
| s(day) |  |  | 118.9 | <0.001 |

We established the lean body mass of males as 142.5 g (SD = 3.06) and females as 148.5 g (SD = 3.93). The mean body mass obtained in first 49 days of the study period was higher than the estimated lean body mass, as in this period average mass of males was 154.0 g (SD = 7.13) and females was 178.0 g (SD = 18.7), with both sexes indicating having stored some fat and protein (Fig. 1S). Predictions for the mass gain made from the best fitted model (Table 1S) indicated an average body mass of 224.3 g for males and 245.6 g for females obtained on last, 91^th^ day of the season (31^st^ August). Assuming average mass from the period of stable body mass as a starting point for mass gain before the departure, males of Great Snipe accumulated 1.63 g per day corresponding to 1.12% of their lean body mass, and females gained 1.57 g per day corresponding to 1.06 % of their lean body mass (Fig. 1S, Table 2S)

## Discussion

Previously obtained results on the African wintering ground of the Great Snipe suggested that both females and males suspend their primary moult prior to departure from their breeding grounds (Debayle *et al*. 2017). Similarly, in our data we found a large proportion of individuals (all males and majority of females) suspending their primary moult, and no birds with active moult detected at the end of the studied period. Moreover, estimated end dates of the primary moult indicate that males should complete the exchange of their primaries in mid-September and females in mid-November. At this time both male and female Great Snipes already arrive at their wintering grounds (Korniluk *et al*. 2015, Lindström *et al*. 2016, Debayle *et al*. 2017), which means that Great Snipes are unable to conclude this process at their breeding grounds. Although the beginning of flight feathers moult on breeding grounds is rare in long distant migratory waders (Kjellén 1994, Remisiewicz 2011), this strategy may be advantageous, as migrants should benefit from expanding their flight feathers moult duration, since slowly grown feathers are of better quality (Dawson *et al*. 2000b, Serra 2001b). Great Snipes may exploit rich feeding sites at their breeding grounds and use resources still available there to partially moult their flight feathers. Than the moult is suspended to avoid migrating with actively growing feathers and with a gap in the wing area, which can generate additional costs for this already demanding process (Swaddle & Witter 1997). Great Snipes resume their flight feathers moult as soon as they reach sub-Saharan Africa in late August, and possibly finish this process before conducting their second, relatively short, intra-African flight. However, Debayle et al. (2017) suggested that some birds may be unable to conclude moult before this intra-African migration and are forced to again suspend their moult or fly with primaries still growing. Therefore advancing the process of flight feathers moult still on breeding grounds might be advantages, as it allows for reducing the amount of time and resources needed for completion of this process on their wintering grounds.

We detected a suspended moult of secondaries only in three males. Moult of secondaries usually starts later than moult of primaries and after reaching a certain threshold of a number of renewed primaries (Summers et al. 2004, Pinchuk & Meissner 2023 personal communication). Indeed, in all three cases birds with signs of started secondary moult were also advanced in their primary moult. It seems however, that moulting of secondaries on their breeding grounds is rather rare in Great Snipes, and majority of individuals conduct this process after reaching wintering grounds.

Female and male Great Snipes have different course of primary moult, with males starting moult earlier in the season and moulting faster compared to females. Moreover, males were characterized by a larger moult range, with a majority of males being able to renew far more primaries before suspending their moult than females. These results are consistent with findings reported by (Debayle *et al*. 2017), where females were less advanced in primary moult than males upon arrival to sub-Sharian Africa. Primary moult end dates given in this study for both sexes are estimated to be approximately one month ahead of dates reported in reality, as on wintering grounds a majority of males finish this process in November, and females in December (Debayle *et al*. 2017). This discrepancy can be explained by the fact that after suspending the moult on breeding grounds both sexes conduct migratory fuelling and later the migratory flight. This creates a temporal delay in the overall completion of moult. The sex-based differences in primary moult timing are most probably caused by the female-only parental care in this lekking species. In June males are still attending leks (Cramp & Simmons 1983), and displaying is an energetically demanding activity (Höglund *et al*. 1992), however, its intensity is possibly lower than in a peak lekking period in May, as suggested by the stable body mass of males at that time (Höglund *et al*. 1992, Witkowska *et al*. 2022). On the contrary in June females are still incubating, rearing the chicks, and investing resources in this process may constrain the initiation of moult. Similar pattern was found in the Purple Sandpiper *Calidris maritima*, where males that are responsible for chick rearing had delayed moult of flight feathers in comparison to females that finish their parental duties after the hatching of eggs (Summers *et al*. 2004). Even slight difference in breeding investment, caused by female producing the clutch may cause delay in the onset of moult. In wader species with parental duties being equally divided between females and males, females begin the process of moult later (Rogers *et al*. 2014, Machín *et al*. 2018).

In females, the standard deviation in the start date of primary moult was relatively large indicating the uncertainty of this estimate. Indeed, in our sample, we noted females with the timing and extent of primary moult resembling those of males, as well as females that did not yet start moulting very late in the studied period, and it is unlikely that they will initiate their moult of flight feathers on breeding grounds. We suggest that females may execute various strategies of primary moult, which possibly can be related to the breeding success of a particular individual. In birds underlying hormonal regulation of moult imposes that maintained breeding activity, causing elevated level of sex-hormones, delays the timing of moult (Dawson 2008). Females with lost broods may moult earlier in the season and at a faster pace, similar to males that are unburdened with parental duties. On the contrary, females with breeding success and therefore large energy expenditure in the chick-rearing period probably moult late in the season, or postpone the moult altogether, until the arrival to the wintering grounds. Indeed, in sub-Saharan Africa, there were some females recorded without any signs of moulting of the flight feathers upon their arrival (Debayle *et al*. 2017). Hence, provided estimates of the primary moult parameters for females based on the Underhill-Zucchini moult model, describe only a general tendency in the primary moult in females.

We found that the scaled body mass index of an individual did not significantly improve the fit of the Underhill-Zucchini moult models, indicating that body condition reflecting the amount of stored energetic resources was not a good predictor for variation in the primary moult parameters. Great Snipes can utilize their fat stores at a quick pace, resulting in large variations in their body mass within a single day (Höglund *et al*. 1992, Witkowska *et al*. 2022). It is possible, that the way of measuring body condition used in this study reflected a current nutritional state of an individual rather than its general quality, albeit a link between a progression of moult and stress caused by malnutrition was shown in other birds (DesRochers *et al*. 2009). Moreover, a negative relationship between moult and the amount of stored energetic resources was described in other bird species (Portugal *et al*. 2007, Alfaro *et al*. 2018), as the overall process of moulting is energetically demanding (Rubolini et al. 2002), with resources used not only for the sole purpose of producing the structure of the feather but also maintaining tissue responsible for the whole process, as well as additional costs related to impaired thermoregulation and flight ability (Buttemer *et al*. 2020). However, larger birds with the slow pace of feather exchange have a relatively low level of daily energetic expenditure devoted to the process of moulting (Lindström *et al*. 1993), that compromise neither flying nor thermoregulation (Hahn *et al*. 1992). Moreover, the lean body mass of birds during the moult changes considerably (Murphy & King 1992, Lind *et al*. 2004), including the ratio of pectoral muscle size to body mass (Lind & Jakobsson 2001), which). Hence, it may explain insignificant influence of the scaled body mass index on primary moult parameters.

In the studied period we detected a non-linear course of fuelling in both female and male Great Snipes. At the beginning of the studied period, birds of both sexes exhibited stable, low scaled mass index reflecting the amount of energetic stores corrected for body size of an individual. This period lasted until the approximately 54^th^ day of the season (25^th^ of July). The stable body mass of lekking males in this period was described in previous studies on the Scandinavian Great Snipe population. This stability, attributed to increased energy intake in late breeding stages, was linked to improved foraging opportunities resulting from snow melt and milder weather conditions (Höglund & Lundberg 1987). However, the lowland population of the Great Snipe do not face such harsh environmental conditions as their Scandinavian conspecifics breeding in the mountains, therefore we think that this explanation is unlikely for birds studied in this work. Stable body condition may be caused by less intense displaying in the late breeding season combined with resource allocation toward the moulting of primary feathers. In females, we also did not find a decrease in those parameters, although chick rearing is considered energetically demanding (Tulp *et al*. 2009, Neubauer *et al*. 2017). We think that suitable conditions on the studied site, ensuring rich feeding sites in a wide range of environmental conditions, create environmental allowing females to maintain the stable body condition, while delaying the onset of their moult (Williams 2018). The period of stable, low body condition overlapped with the time of the moult for males Great Snipe and moult and/or chick rearing of females. The last birds, (two females and one male) with active moult were detected in our sample on the 58^th^ day of the season (29^th^ of July). Moreover, as most males suspend their moult on the 5^th^ or 6^th^ primary, and females on their 1^st^ or 2^nd^ primary, we could predict based on the modelled duration that the majority of males should suspend their moulting on breeding grounds on the 55^nd^ to 64^st^ day of the season (27^th^ July and 4^th^ August respectively) and females should do so from 55^th^ to 62^th^ day of the season (26^th^ July and 2^nd^ of August). This indicates that moulting and fuelling for migration do not overlap or overlap to a small extent, and indicates the trade-off in resource allocation between those two processes.

In the studied period both females and males had similar energetic resources stored, but significant interaction between sex and day of the season in the changes of the scaled mass index over time, indicates that males increase their energetic stores at a faster pace compared to females. Males end their breeding performance before females, as they do not participate in chick rearing, and are able to finish their moult sooner, despite having a larger range of exchanged primaries. This leads to effectively more time for males to exploit resources and fuel for migration, which could account for this effect. Moreover, males arrive sooner to sub-Saharan Africa compared to females, explaining their need for more rapid fuelling (Debayle *et al*. 2017). The overall rate of fuelling and maximum mass obtained in both sexes was similar to mass gain rates reported for fuelling on wintering grounds, before northward migratory flight (Debayle *et al*. 2017). Still, comparing the fuel load of other waders of a similar size (Gudmundsson *et al*. 1991, Kvist & Lindström 2003, Piersma *et al*. 2005), the fuelling rate presented in our study is relatively low for a long-distance migrant, covering 5000 km with a non-stop flight (Lindström *et al*. 2016). As Great Snipes begin moult of their wing feathers on breeding grounds prior to fuelling and as there is some plasticity in the extent of moult between individuals, it seems that gathering energetic stores for flight is not greatly time constrained. It is possible that we failed to detect fat Great Snipes ready for departure, although this would be the second attempt to describe the process of fuelling for migratory flight in this species that would do so (Debayle *et al*. 2017). Great Snipes reach Sub-Saharan Africa in late September (Debayle *et al*. 2017), and the last birds that we were able to measure in this study were caught on 1^st^ of September. Time necessary for conducting trans-Sahara flight takes up to 3 days of non-stop flight (on average 64 hours) for Great Snipe males of Scandinavian population (Lindström *et al*. 2016). Therefore, there is still time left for Great Snipes to accumulate sufficient energetic stores before departure from European breeding grounds, especially that birds might increase their energy intake for more rapid migratory fuelling before departure (Meissner *et al*. 2011, Lindström *et al*. 2019).

## Conclusion

In this study, we examined the primary moulting and fuelling strategies of Great Snipes, highlighting the balance between the reproductive efforts, moult, and migratory preparation for this species. Our research supported previous findings from wintering grounds, indicating that both females and males of this species suspend their primary moult before departing from their European breeding grounds. Males initiate primary moult earlier, moult faster resulting in a larger moult range of renewed primaries, compared to females. The intersexual differences in primary moult strategies stem from distinct parental roles within this lekking species. Females, responsible for incubation and chick rearing, might delay their moult, whereas males, unburdened with parental duties could begin moulting their flight feathers earlier. We observed a non-linear pattern of fuelling in both female and male Great Snipes. Stable and low body condition was maintained until late July overlapping with moult and/or chick-rearing period, suggesting a trade-off in resource allocation between these processes. Later in the season body condition increased, however, the fuelling rate of 1% of the lean body mass increase per day observed in Great Snipe appeared relatively low for a species undertaking a long-distance migration. It is possible that the accumulation rate of fat stores increases towards departure, similar to other wader species.

## Acknowlegments

We are grateful to all the volunteers who worked at the Turov Ringing Station for the past 23 years, helping in gathering data used in this study, in particular A. Zyatikov, I. Bogdanovich, A. Usau, V. Khursanau, A. Khalandach and I. Kashpei.

## Authors contribution

MW: Conceptualization, data curation, formal analysis, investigation, methodology, visualization, writing – original draft, writing – review and editing

MK: Conceptualization, data curation, investigation, methodology, writing – review and editing PP: Conceptualization, data curation, investigation, methodology, writing – review and editing

TT: Data curation, investigation, writing – review and editing

NK: Data curation, investigation, writing – review and editing

WM: Conceptualization, data curation, investigation, methodology, writing – review and editing

## Ethical note

All conducted procedures were in accordance with Belarussian and Polish law.

## Funding

This study was possible thanks to funding from the University of Gdańsk, the National Academy of Sciences of Belarus, and APB BirdLife Belarus. Part of the fieldwork caried out in Poland was funded by the EU LIFE Programme under the project ‘Implementation of the National Action Plan for Great Snipe in Poland – phase I’ LIFE17 NAT/PL/000015 and the National Fund for Environment Protection and Water Management in Warsaw, coordinated by Lubelskie Towarzystwo Ornitologiczne (LTO) and Natura International Polska

## Conflict of interest

The authors declare no conflict of interest.

## Data availability statement

Data and code are available at the repository RepOD: (https://repod.icm.edu.pl/privateurl.xhtml?token=845a0c53-c009-4d51-bca3-87c9bea56df8), doi: https://doi.org/10.18150/LB46RA (DOI is inactive before publication of this work)

## Notes

### Competing Interest Statement

The authors have declared no competing interest.

## References

Alerstam, T. & Lindström, Å. 1990. Optimal Bird Migration: The Relative Importance of Time, Energy, and Safety. In: Bird Migration, pp. 331–351. Springer-Verlag Berlin, Heidelberg.

Alfaro, M., Sandercock, B.K., Linguori, L. & Arim, M. 2018. Body condition and feather molt of a migratory shorebird during the non-breeding season. J Avian Biol 49.

Bartoń, K. 2023. MuMIn: Multimodal Inference. R package ver. 1.47.5.

Bonier, F., Martin, P.R., Jensen, J.P., Butler, L.K., Ramenofsky, M. & Wingfield, J.C. 2007. Pre-migratory life history stages of juvenile arctic birds: Costs, constraints, and trade-offs. Ecology 88: 2729–2735.

Burnham, K.P. & Anderson, D.R. 2004. Multimodel Inference: Understanding AIC and BIC in Model Selection. Sociol Methods Res 33: 261–304.

Busse, P. & Meissner, W. 2015. Bird ringing station manual. De Gruyer, Warsaw/Berlin.

Buttemer, W.A., Addison, B.A. & Klasing, K.C. 2020. The energy cost of feather replacement is not intrinsically inefficient. Can J Zool 98: 142–148.

Conklin, J.R., Battley, P.F. & Potter, M.A. 2013. Absolute Consistency: Individual versus Population Variation in Annual-Cycle Schedules of a Long-Distance Migrant Bird. PLoS One 8.

Cramp, S. & Simmons, K.E.L. 1983. The birds of the western Palearctic,vol III. Oxford University Press, Oxford. Oxford University Press, Oxford.

Dawson, A. 2008. Control of the annual cycle in birds: Endocrine constraints and plasticity in response to ecological variability. Philosophical Transactions of the Royal Society B: Biological Sciences 363: 1621–1633.

Dawson, A., Hinsley, S.A., Ferns, P.N., Bonser, R.H.C. & Eccleston, L. 2000a. Rate of moult affects feather quality: A mechanism linking current reproductive effort to future survival. Proceedings of the Royal Society B: Biological Sciences 267: 2093–2098.

Dawson, A., Hinsley, S.A., Ferns, P.N., Bonser, R.H.C. & Eccleston, L. 2000b. Rate of moult affects feather quality: A mechanism linking current reproductive effort to future survival. Proceedings of the Royal Society B: Biological Sciences 267: 2093–2098.

Debayle, E.J.M., Devort, M., Klaassen, R.H.G. & Lindström, Å. 2017. Great Snipes in sub-Saharan Africa: Seasonal patterns of abundance, moult and body mass in relation to age and sex. Wader Study 124: 186–196.

DesRochers, D.W., Reed, J.M., Awerman, J., Kluge, J.A., Wilkinson, J., van Griethuijsen, L.I., Aman, J. & Romero, L.M. 2009. Exogenous and endogenous corticosterone alter feather quality. Comparative Biochemistry and Physiology - A Molecular and Integrative Physiology 152: 46–52.

Erni, B., Bonnevie, B.T., Oschadleus, H.D., Altwegg, R. & Underhill, L.G. 2013. Moult: An R package to analyze moult in birds. J Stat Softw 52: 1–23.

Faaborq, J., Holmes, R.T., Anders, A.D., Bildstein, K.L., Dugger, K.M., Gauthreaux, S.A., Heglund, P., Hobson, K.A., Jahn, A.E., Johnson, D.H., Latta, S.C., Levey, D.J., Marra, P.P., Merkord, C.L., Erica, N.O.L., Rothstein, S.I., Sherry, T.W., Scott Sillett, T., Thompson, F.R. & Warnock, N. 2010. Recent advances in understanding migration systems of New World land birds. Ecol Monogr 80: 3–48.

Ginn, H. & Melville, D.S. 1983. Moult in birds. British Trust for Ornithology, Tring.

Gudmundsson, G.A., Lindstrom, A.K.E. & Alerstam, T. 1991. Optimal fat loads and long-distance flights by migrating Knots Calidris canutus, Sanderlings C. alba and Turnstones Arenaria interpres. Ibis 133: 140–152.

Hahn, T.P., Swingle, J., Wingfield, J.C. & Ramenofsky, M. 1992. Adjustments of the prebasic molt schedule in birds. Ornis Scandinavica 23: 314–321.

Hastie, T. & Tibsgirani, R. 1986. Additive Models Generalized. Statistical Science 1: 297–310.

Helm, B. & Gwinner, E. 2006. Timing of molt as a buffer in the avian annual cycle. Acta Zoologica Sinica 52: 703–706.

Hemborg, C. & Lundberg, A. 1998. Costs of overlapping reproduction and moult in passerine birds: An experiment with the pied flycatcher. Behav Ecol Sociobiol 43: 19–23.

Höglund, J., Kålås, J.A. & Fiske, P. 1992. The costs of secondary sexual characters in the lekking great snipe (Gallinago media). Behav Ecol Sociobiol 30: 309–315.

Hoglund, J., Kalas, J.A. & Lofaldli, L. 1990. Sexual dimorphism in the lekking great snipe.

Höglund, J. & Lundberg, A. 1987. Sexual selection in a monomorphic lek-breeding bird: correlates of male mating success in the great snipe Gallinago media. Behav Ecol Sociobiol 21: 211–216.

Kjellén, N. 1994. Moult in relation to migration in birds - a review. Ornis Svec 4: 1–24.

Korniluk, M., Białomyzy, P., Grygoruk, G., Kozub, Ł., Sielezniew, M., Świętochowski, P., Tumiel, T., Wereszczuk, M. & Chylarecki, P. 2020. Habitat selection of foraging male Great Snipes on floodplain meadows: importance of proximity to the lek, vegetation cover and bare ground. Ibis, doi: 10.1111/ibi.12898.

Korniluk, M. & Chylarecki, P. 2023. Intra-Seasonal Lek Changes of Great Snipe Gallinago media Males in the Northeast of Poland. Acta Ornithol 58.

Korniluk, M., Tumiel, T., Świętochowski, P., Wereszczuk, M., Białomyzy, P., Grygoruk, G. & Iliszko, L. 2015. Migration pattern and behaviour of the Great Snipe Gallinago media lowland population. In: *International Wader Study Group Conference*. Ásbrú.

Kvist, A. & Lindström, Å. 2003. Gluttony in migratory waders - Unprecedented energy assimilation rates in vertebrates. Oikos 103: 397–402.

Labocha, M.K. & Hayes, J.P. 2012. Morphometric indices of body condition in birds: A review. J Ornithol 153: 1–22.

Lind, J., Gustin, M. & Sorace, A. 2004. Compensatory bodily changes during moult in Tree Sparrows Passer montanus in Italy. Ornis Fenn 81: 75–83.

Lind, J. & Jakobsson, S. 2001. Body building and concurrent mass loss: Flight adaptations in tree sparrows. Proceedings of the Royal Society B: Biological Sciences 268: 1915–1919.

Lindström, Å. 2003. Fuel Deposition Rates in Migrating Birds: Causes, Constraints and Consequences. In: Avian Migration (P. Berthold, E. Gwinner, & E. Sonnenschein, eds). Springer, Berlin, Heidelberg.

Lindström, Å., Alerstam, T., Andersson, A., Bäckman, J., Bahlenberg, P., Bom, R., Ekblom, R., Klaassen, R.H.G., Korniluk, M., Sjöberg, S. & Weber, J.K.M. 2021. Extreme altitude changes between night and day during marathon flights of great snipes. Current Biology 31: 3433–3439.e3.

Lindström, Å., Alerstam, T., Bahlenberg, P., Ekblom, R., Fox, J.W., Råghall, J. & Klaassen, R.H.G. 2016. The migration of the great snipe Gallinago media: Intriguing variations on a grand theme. J Avian Biol 47: 321–334.

Lindström, Å., Alerstam, T. & Hedenström, A. 2019. Faster fuelling is the key to faster migration. Nat Clim Chang 9: 288–289.

Lindström, Å., Daan, S. & Visser, G.H. 1994.The conflict between moult and migratory fat deposition: A photoperiodic experiment with bluethroats.

Lindström, Å., Visser, G.H. & Daan, S. 1993. The Energetic Cost of Feather Synthesis Is Proportional to Basal Metabolic Rate. Physiol Zool 66: 490–510.

Machín, P., Remisiewicz, M., Fernández-Elipe, J., Jukema, J. & Klaassen, R.H.G. 2018. Conditions at the breeding grounds and migration strategy shape different moult patterns of two populations of Eurasian golden plover Pluvialis apricaria. J Avian Biol 49: 1–12.

Mazur, A.E., Remisiewicz, M. & Underhill, L.G. 2021. Sex-specific patterns of fuelling and pre-breeding body moult of Little Stints Calidris minuta in South Africa. Ibis 163: 99–112.

McNamara, J.M., Welham, R.K. & Houston, A.I. 1998. The Timing of Migration within the Context of an Annual Routine. J Avian Biol 29: 416–423.

Meissner, W., Karlionova, N. & Pinchuk, P. 2011. Fuelling Rates by Spring-Staging Ruffs Philomachus pugnax in Southern Belarus. Ardea 99: 147–155.

Meissner, W., Zaniewicz, G., Gogga, P., Pilacka, L., Klaassen, M. & Minton, C. 2018. Relative mass of flight feathers in waders – an update. Wader Study 125: 205–211.

Morrison, C.A., Alves, J.A., Gunnarsson, T.G., Þórisson, B. & Gill, J.A. 2019. Why do earlier-arriving migratory birds have better breeding success? Ecol Evol 9: 8856–8864.

Murphy, M.E. & King, J.R. 1992. Energy and nutrient use during moult by White-crowned Sparrows Zonotrichia gambelii leucophrys. Ornis Scandinavica 23: 304–313.

Neubauer, G., Pilacka, L., Zieliński, P. & Gromadzka, J. 2017. Population-level body condition correlates with productivity in an arctic wader, the dunlin Calidris alpina, during post-breeding migration. PLoS One 12: 1–17.

Peig, J. & Green, A.J. 2009. New perspectives for estimating body condition from mass/length data: The scaled mass index as an alternative method. Oikos 118: 1883–1891.

Piersma, T., Rogers, D., González, P.M., Zwarts, L., Niles, L.J., Lima, I., Donascimento, S., Minton, C.D.T. & Baker, A. 2005. Fuel storage rates before northward flights in Red Knots worldwide - Facing the Severest Ecological Constraint in Tropical Intertidal Environments? In: Birds of two worlds: the ecology and evolution of migratory bird. (R. Greenberg & P. P. Marra, eds), pp. 262–274. The Johns Hopkins University Press, Baltimore, London.

Pinchuk, P. V., Karlionova, N. V., Bogdanovich, I.A., Luchik, E.A. & Meissner, W. 2016. Age and seasonal differences in biometrics of dunlin (Calidris Alpina) migrating in spring through the pripyat river floodplain, Southern Belarus. Zool Zhurnal 95: 327–334.

Portugal, S.J., Green, J.A. & Butler, P.J. 2007. Annual changes in body mass and resting metabolism in captive barnacle geese (Branta leucopsis): The importance of wing moult. Journal of Experimental Biology 210: 1391–1397.

R Core Team. 2022. R: A language and environment for statistical computing.

Remisiewicz, M. 2011. The flexibility of primary moult in relation to migration in Palaearctic waders - An overview. Wader Study Group Bulletin 118: 163–174.

Ricklefs, R.E. 1996. Avian Energetics, Ecology, and Evolution. In: Avian Energetics and Nutritional Ecology. Springer, Boston.

Rogers, K.G., Rogers, D.I. & Weston, M.A. 2014. Prolonged and flexible primary moult overlaps extensively with breeding in beach-nesting Hooded Plovers Thinornis rubricollis. Ibis 156: 840– 849.

Rubolini, D., Massi, A. & Spina, F. 2002. Replacement of body feathers is associated with low pre-migratory energy stores in a long-distance migratory bird, the barn swallow (Hirundo rustica). J Zool 258: 441–447.

Serra, L. 2001a. Duration of primary moult affects primary quality in Grey Plovers Pluvialis squatarola. J Avian Biol 32: 377–380.

Serra, L. 2001b. Duration of primary moult affects primary quality in Grey Plovers Pluvialis squatarola. J Avian Biol 32: 377–380.

Stutchbury, B.J.M., Gow, E.A., Done, T., MacPherson, M., Fox, J.W. & Afanasyev, V. 2011. Effects of post-breeding moult and energetic condition on timing of songbird migration into the tropics. Proceedings of the Royal Society B: Biological Sciences 278: 131–137.

Summers, R.W., Underhill, L.G., Nicoll, M., Strann, K.B. & Nilsen, S. 2004. Timing and duration of moult in three populations of Purple Sandpipers Calidris maritima with different moult/migration patterns. Ibis 146: 394–403.

Swaddle, J.P. & Witter, M.S. 1997. The effects of molt on the flight performance, body mass, and behavior of European starlings (Sturnus vulgaris): An experimental approach. Can J Zool 75: 1135–1146.

Terrill, R.S. & Shultz, A.J. 2022. Feather function and the evolution of birds. Biological Reviews 98: 540–566.

Tulp, I., Schekkerman, H., Bruinzeel, L.W., Jukema, J., Visser, G.H. & Piersma, T. 2009. Energetic demands during incubation and chick rearing in a uniparental and a biparental shorebird breeding in the high arctic. Auk 126: 155–164.

Underhill, L.G. & Zucchini, W. 1988. A model for avian primary moult. Ibis 130: 358–372.

Underhill, L.G., Zucchini, W. & Summers, R.W. 1990. A model for avian primary moult-data types based on migration strategies and an example using the Redshank Tringa totanus. Ibis 132: 118– 123.

Vágási, C.I., Pap, P.L., Vincze, O., Benko, Z., Marton, A. & Barta, Z. 2012. Haste makes waste but condition matters: Molt rate-feather quality trade-off in a sedentary songbird. PLoS One 7: 1–10.

Williams, T.D. 2018. Physiology, activity and costs of parental care in birds. Journal of Experimental Biology 221.

Wingfield, J.C. 2008. Organization of vertebrate annual cycles: Implications for control mechanisms. Philosophical Transactions of the Royal Society B 363: 425–441.

Witkowska, M., Pinchuk, P., Meissner, W. & Karlionova, N. 2023. Body size constrains the annual apparent survival of lekking Great Snipe Gallinago media males of eastern, lowland population. J Ornithol, doi: 10.1007/s10336-023-02091-7.

Witkowska, M., Pinchuk, P., Meissner, W., Karlionova, N. & Marynkiewicz, Z. 2022. The level of water in the river flowing through the breeding site shapes the body condition of a lekking bird—the Great Snipe Gallinago media. J Ornithol 163: 385–394.

Wood, A.S., Scheipl, F. & Wood, M.S. 2020. Package ‘gamm4’.

