## Supplementary Online Material for "The interplay between moult of flight feathers and fuelling conducted on the breeding grounds of the Great Snipe *Gallinago media* from the eastern European, lowland population"

[3] – Nature Association Dubelt, Juszkowy Gród 17, Michałowo, 16-050, Poland

[4] – APB BirdLife Belarus, Minsk, Belarus

[5] – National Park 'Pripyatsky', Turov, Belarus

[6] – Scientific and Practical Center for Biological Resources, National Academy of Sciences of Belarus, Minsk, Belarus

ORCID

Marta Witkowska: 0000-0001-8153-0270

Michał Korniluk: 0000-0002-4367-093

Natalia Karlionova: 0000-0002-3266-7251

Włodzimierz Meissner: 0000-0001-5995-9185

In this work to analyse the process of fuelling before departure, we decided to additionally present the second set of models using body mass instead of the scaled mass index as a dependent variable. We included a null model, with intercept as the sole factor, the global model with the day of the season as a smooth term, sex, and its interaction with the day of the season as independent variables. We also established two reduced, nested models: a model with the day of the season as a single independent variable, and a model with the day of the season as a smooth term and the sex of an individual as an independent variable. Model selection was carried out based on the Akaike Information Criterion and Akaike weights. In the case of models with delta AICc lower than 2, indicating their similar parsimony the estimates were derived from full model averaging (Burnham & Anderson, 2004). Model fit was also described using $R^2$ and the percentage of explained deviance. The models were fit using *gamm4* package (Wood et al., 2020), with model selection and averaging performed using *MuMIn* package (Bartoń, 2023) in R version 4.2.2 (R Core Team, 2022)

Ranking of GAM models used to describe fuelling before departure revealed similar results in both sets of models with either body mass or scaled mass index as a depended variable. In case of the first set of models, two top ranking models were distinguished as similarly parsimonious based on their ΔAICc < 2 and approximate values of $w_i$, as well as having similar fit to the data (models BM1 and BM2, Table 1S). These models included day of the season as a smooth term, sex and/or interaction between sex and day of the season. Averaging their results indicated that the day of the season as smooth term significantly influenced body mass, where this parameter was stable till the approximately 49th day of the season (20th July) with later increase (Fig. 1S, Table 2S). Moreover, females had significantly higher body mass compared to males, with statistically insignificant interaction between sex and day of the season as an independent variable (Table 2S).

Table 1S. Ranking of GAMs estimating the relationship of body mass with sex and day of the season. 1 in model formula indicates null model with only intercept included as an independent variable. s() – independent variable used as a smooth factor, + - additive relationship, * - interaction between variables, edf – effective degrees of freedom, AICc – Akaike's Information Criterion for small sample size, ΔAICc – difference in AICc between the given model and model with the lowest AICc value, $w_i$ – Akaike weight. Adjusted $R^2$ and percentage of explained deviance is given. Top ranking models with the ΔAICc < 2 are bolded.

| Model ID | Model formula | edf | AICc | ΔAICc | $w_i$ | adj. $R^2$ | % Dev. Exp. |
|---|---|---|---|---|---|---|---|
| A) Body mass as a dependent variable | | | | | | | |
| **BM1** | **BM = s(day) + sex** | **9.13** | **2142.77** | **0** | **0.579** | **0.692** | **70.0%** |
| **BM2 (global)** | **BM = s(day) + sex + sex*day** | **10.08** | **2143.41** | **0.64** | **0.421** | **0.632** | **70.2%** |
| BM3 | BM = s(day) | 8.22 | 2463.71 | 320.94 | < 0.001 | 0.568 | 57.7% |
| BM4 (null) | BM = 1 | 2 | 2705.30 | 562.53 | < 0.001 | 0.000 | 0% |

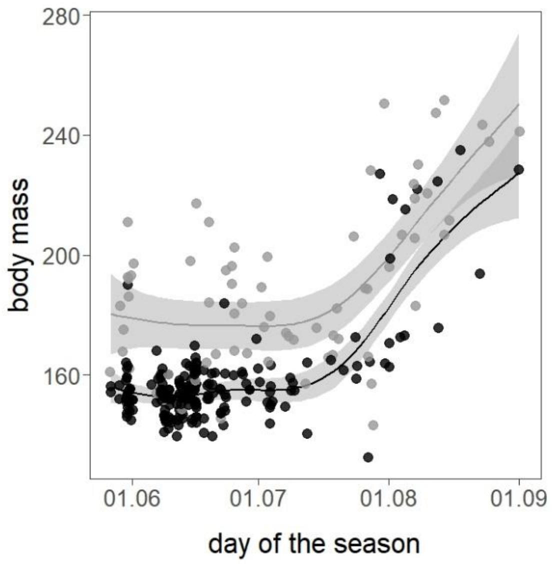


Fig. 1S. Changes of body mass during the season in adult females (grey) and males (black). Line – significant relationship estimated with the Generalized Additive Model, grey area – 95% confidence interval.

Table 2S. Results of top ranking GAMs (BM1 i BM2 averaged) explaining the relationship between body mass and sex, interaction between sex and the day of the season and day of the season as a smooth factor.

| Fixed effect | Estimate | SE | *z* | *P* |
|---|---|---|---|---|
| intercept | 134.67 | 56.85 | 2.37 | 0.018 |
| sex (M) | -22.72 | 3.19 | 7.10 | <0.001 |
| sex (M) * day | 0.04 | 0.07 | 0.56 | 0.57 |
| Smoothed term | | | *F* | *P* |
| s(day) | | | 60.51 | <0.001 |

Overall using body mass instead of scaled mass index for describing the process of fuelling for the migratory flight in Great Snipes gave similar results. The two differences reported are:

- Sex was a significant factor influencing body mass, but not scaled mass index. This is caused by the sexual dimorphism in size in Great Snipes, where females are the bigger and, therefore heavier sex (Hoglund et al., 1990).
- Males increase their energetic stores at a faster pace compared to females, although this effect was detected only in model using the scaled mass index as a dependent variable. This is possibly due to the presence of one small male, with relatively small body mass caught late in the season, that resulted in insignificant interaction between sex and day of the season. We decided to not treat this record as an outlier, as this individual gathered sufficient energetic stores for this period, when its body mass was corrected by its body size.

Literature:

Bartoń, K. (2023). MuMIn: Multimodal Inference. *R Package Ver. 1.47.5*.

Burnham, K. P., & Anderson, D. R. (2004). Multimodel Inference: Understanding AIC and BIC in Model Selection. *Sociological Methods and Research*, *33*, 261–304.

Hoglund, J., Kalas, J. A., & Lofaldli, L. (1990). Sexual dimorphism in the lekking great snipe. In *Ornis Scandinavica* (Vol. 21, Issue 1, pp. 1–6). https://doi.org/10.2307/3676372

R Core Team. (2022). *R: A language and environment for statistical computing.* R Foundation for Statistical Computing. https://www.r-project.org/

Wood, A. S., Scheipl, F., & Wood, M. S. (2020). *Package 'gamm4'* (version 0.2-6). R package. https://cran.r-project.org/package=gamm4
